# TIGA: Target illumination GWAS analytics

**DOI:** 10.1101/2020.11.11.378596

**Authors:** Jeremy J Yang, Dhouha Grissa, Christophe G Lambert, Cristian G Bologa, Stephen L Mathias, Anna Waller, David J Wild, Lars Juhl Jensen, Tudor I Oprea

**Affiliations:** Department of Internal Medicine, Division of Translational Informatics, University of New Mexico Health Sciences Center, Albuquerque, NM, USA; Department of Pathology, University of New Mexico Health Sciences Center, Albuquerque, NM, USA; Novo Nordisk Foundation Center for Protein Research, Faculty of Health and Medical Sciences, University of Copenhagen, Copenhagen, Denmark; School of Informatics, Computing and Engineering, Indiana University, Bloomington, IN, USA

**Keywords:** GWAS, data science, drug discovery, drug target, druggable genome

## Abstract

Genome wide association studies (GWAS) can reveal important genotype–phenotype associations, however, data quality and interpretability issues must be addressed. For drug discovery scientists seeking to prioritize targets based on the available evidence, these issues go beyond the single study. Here, we describe rational ranking, filtering and interpretation of inferred gene–trait associations and data aggregation across studies by leveraging existing curation and harmonization efforts. Each gene–trait association is evaluated for confidence, with scores derived solely from aggregated statistics, linking a protein-coding gene and phenotype. We propose a method for assessing confidence in gene–trait associations from evidence aggregated across studies, including a bibliometric assessment of scientific consensus based on the iCite Relative Citation Ratio, and meanRank scores, to aggregate multivariate evidence. This method, intended for drug target hypothesis generation, scoring and ranking, has been implemented as an analytical pipeline, available as open source, with public datasets of results, and a web application designed for usability by drug discovery scientists, at https://unmtid-shinyapps.net/tiga/.

## Introduction

Over the two decades since the first draft human genome was published, dramatic progress has been achieved in foundational biology with translational benefits to medicine and human health. Genome wide association studies (GWAS) contribute to this progress by inferring associations between genomic variations and phenotypic traits (Bossé and Amos, 2018; Rusu *et al*., 2017). These associations are correlations which may or may not be causal. While GWAS can reveal important genotype–phenotype associations, data quality and interpretability must be addressed (Lambert and Black, 2012; Visscher *et al*., 2017; Marigorta *et al*., 2018; Gallagher and Chen-Plotkin, 2018). For drug discovery scientists seeking to prioritize targets based on evidence from multiple studies, quality and interpretability issues are broader than for GWAS specialists. For this use case, GWAS are one of several evidence sources to be explored and considered, and interpretability must be in terms of genes corresponding to plausible targets, and traits corresponding to diseases of interest.

Single nucleotide variants (SNV) are the fundamental unit of genomic variation, and the term single nucleotide polymorphism (SNP) refers to SNVs identified as common sites of variation relative to a reference genome, and measured by microarray or sequencing technologies. The NHGRI-EBI GWAS Catalog (Buniello *et al*., 2019) -- hereafter “Catalog” – curates associations between SNPs and traits from GWAS publications, shares metadata and summary data, standardizes heterogeneous submissions, maps formats and harmonizes content, mitigating widespread data and meta-data issues according to FAIR (Findable, Accessible, Interoperable and Reusable) principles (Wilkinson *et al*., 2016). These challenges are exacerbated by rapid advances in experimental and computational methodology. As *de facto* GWAS registrar, the Catalog interacts directly with investigators and accepts submissions of summary statistic data in advance of publication. Proposing and maintaining metadata standards the Catalog advocates and advances FAIRness in GWAS, for the benefit of the community. The Catalog addresses many difficulties due to content and format heterogeneity, but there are continuing difficulties and limitations both from lack of reporting standards and the variability of experimental methodology and diagnostic criteria.

Other GWAS data collections include the Genome-Wide Repository of Associations between SNPs and Phenotypes, GRASP (Eicher *et al*., 2015) and The Framingham Heart Study, which employs non-standard phenotypes and some content from the Catalog (not updated since 2015). GWASdb (Li *et al*., 2016) integrates over 40 data sources in addition to the Catalog, includes less significant variants to address a variety of use cases, and has been maintained continually since 2011. GWAS Central, continually updated through 2019, includes less significant associations and provides tools for a variety of exploration modes based on Catalog data, but is not freely available for download. PheGenI (Ramos *et al*., 2014) integrates Catalog data with other NCBI datasets and tools. Others integrate GWAS with additional data (e.g. pathways, expression, linkage disequilibrium) to associate traits or diseases with genes (Greene *et al*., 2015; Shen *et al*., 2017; Wainberg *et al*., 2019; Li *et al*., 2018; Pallejà *et al*., 2012). Each of these resources offers unique value and features. For this use case, the Catalog is the logical choice, given its applicability and commitment to expert curation, data standards, support and maintenance.

Here we describe TIGA (Target Illumination GWAS Analytics), an application for illuminating understudied drug targets. TIGA enables ranking, filtering and interpretation of inferred gene-trait associations aggregated across studies from the Catalog. Each inferred gene-to-trait association is evaluated for confidence, with scores derived solely from evidence aggregated across studies, linking a phenotypic trait and protein-coding gene, mapped from single nucleotide polymorphism (SNP) variation. TIGA uses the Relative Citation Ratio, RCR (Hutchins *et al*., 2016), a bibliometric statistic from iCite (Hutchins *et al*., 2019). TIGA does not index the full corpus of GWAS associations, but focuses on the strongest associations at the protein-coding gene level instead, filtered by disease areas that are relevant to drug discovery. For instance, GWAS for highly polygenic traits are considered less likely to illuminate druggable genes. Here, we describe the web application and its interpretability for non-GWAS specialists. We discuss TIGA as an application of data science for scientific consensus and interpretability, including statistical and semantical challenges. Code and data are available under BSD-2-Clause license from https://github.com/unmtransinfo/tiga-gwas-explorer.

## Methods

### NHGRI-EBI GWAS Catalog preprocessing

The 2021-02-12 release of the Catalog references 11671 studies and 4865 PubMed IDs. The curated associations include 8235 studies and 2706 EFO-mapped traits. After filtering studies to require (1) mapped trait, (2) p-value below 5e-8, (3) reported effect size (odds-ratio or beta), and (4) mapped protein-coding gene, we obtain 4118 studies, 1521 traits, and 10264 genes. For consistency between studies, only genes mapped by the Ensembl pipeline for genomics annotations^1^ were considered (not author-reported). Figures 1 and 2 illustrate the growth of GWAS research as measured by counts of studies and subjects.

**Fig 1:**
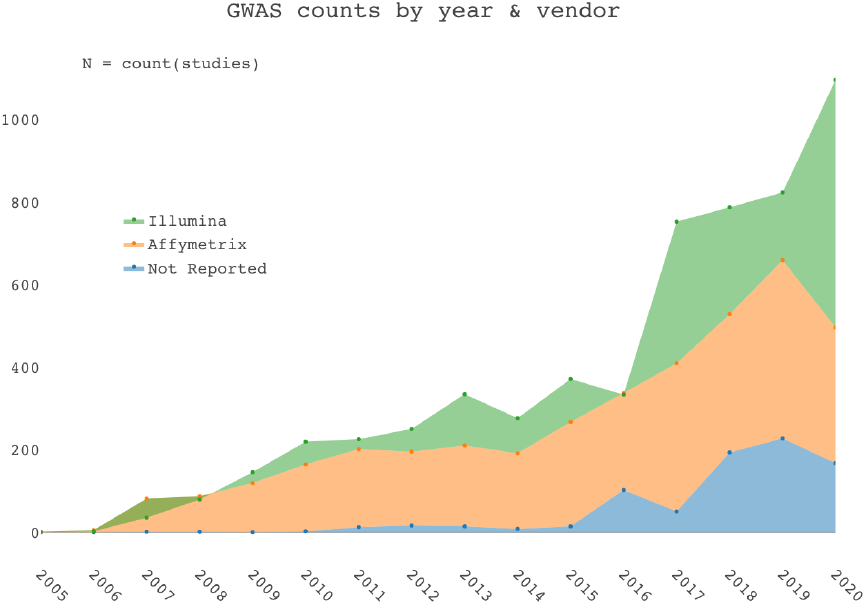
GWAS Catalog study counts by year and vendor, indicating growth and platform trends.

**Fig 2:**
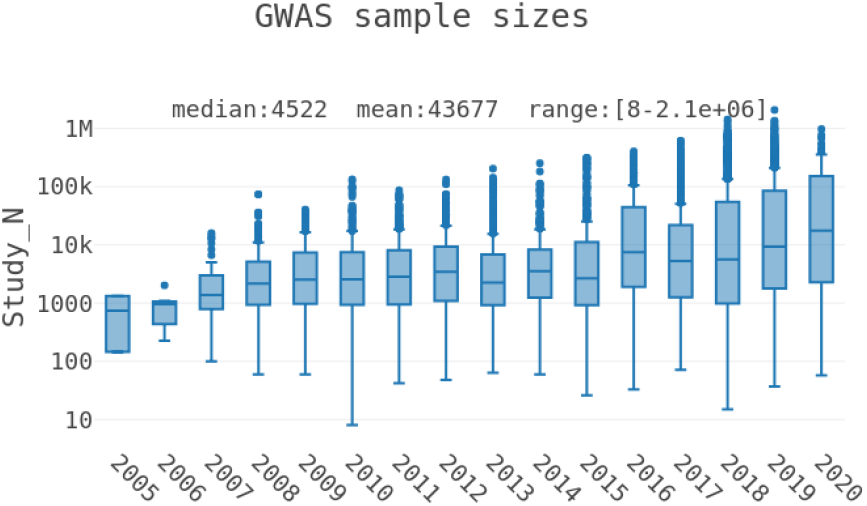
GWAS Catalog sample size distributions by year, on log scale, indicating variance in statistical power.

### RCRAS = Relative Citation Ratio (RCR) Aggregated Score

The purpose of TIGA is to evaluate the evidence for a gene-trait association, by aggregating multiple studies ***and*** their corresponding publications. The iCite RCR (Hutchins *et al*., 2016) is a bibliometric statistic designed to evaluate the impact of an individual publication (in contrast to the journal impact factor). By field- and time-normalizing per-publication citation counts, the RCR measures evolving impact, in effect a proxy for scientific consensus. Hence by aggregating RCRs we seek a corresponding measure of scientific consensus for associations.

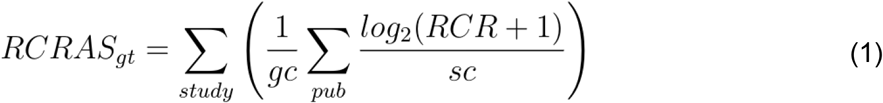

Where:

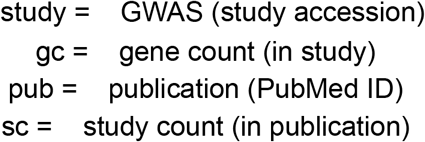

The log_2_() function is used with the assertion that differences of evidence depend on relative, rather than absolute differences in RCR. Division by sc effects a partial count for publications associated with multiple studies. Since RCR≥0, log_2_(RCR+1)≥ 0 and intuitively, when RCR= 1 and sc= 1, log_2_(RCR+ 1) = 1. Similarly division by gc reflects a partial count since studies may implicate multiple genes. This approach is informed by bibliometric methodology, including fractional counting, as described elsewhere (Cannon *et al*., 2017). For recent publications lacking RCR, we used the global median as an estimated prior. Computed thus, RCRAS extends RCR with similar logic, providing a rational bibliometric measure of evidence for scoring and ranking gene-trait associations.

### Association weighting by SNP–gene distance

Mapping genomic variation of single nucleotides (SNPs) to genes is a challenging area of active research (Liu *et al*., 2010; Mishra and Macgregor, 2015; Lamparter *et al*., 2016). This project does not contribute to mapping methodology. Rather, TIGA employs mappings provided by the Catalog between GWAS SNPs and genes, generated by an Ensembl pipeline, which “adds additional SNP specific information associated with the rsID extracted… This information is retrieved using the Ensembl API and the source of the data is both Ensembl and NCBI.”(The NHGRI-EBI GWAS Catalog) It is important to note that this method is unbiased and derived from experimental data and the current human reference genome. TIGA aggregates SNP-trait associations, assessing evidence for gene-trait associations, based on these understandings:

- SNPs within a gene are more strongly associated than SNPs upstream or downstream.
- Strength of association decreases with distance, or more rigorously stated, the probability of linkage disequilibrium (LD) between a SNP and protein coding gene decreases with genomic physical distance. Accordingly, we employ an inverse exponential scoring function, consistent with LD measure (**Δ**) and coefficient of decay (**β**) by Wang and coworkers (Wang *et al*., 2006).

This function, used to weight N_snp to compute a distance-weighted SNP count N_snpw, is plotted together with the observed frequencies of mapped gene distances in supplementary Fig. 1, to illustrate how the extant evidence is weighted.

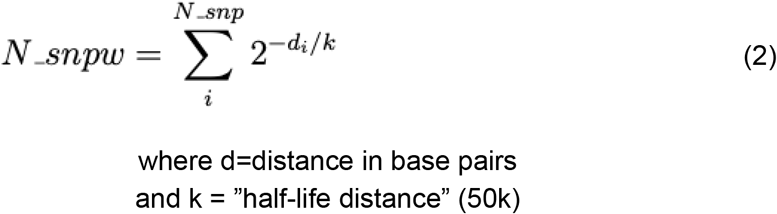

### Multivariate ranking

Multivariate ranking is a well studied problem which needs to be addressed for ranking GWAS associations. We evaluated two approaches, namely non-parametric **μ** scores (Wittkowski, 2008) and meanRank, and chose the latter based on benchmark test performance. meanRank aggregates ranks instead of variables directly, avoiding the need for *ad hoc* parameters. Variable-ties imply rank-ties, with missing data ranked last. We normalize scoring to (0,100] defining **meanRankScore** as follows.

Variables of merit used for scoring and ranking gene-trait associations:

- N_snpw: N_snp weighted by distance inverse exponential described above.
- pVal_mLog: max(-Log(pValue)) supporting gene-trait association.
- RCRAS: Relative Citation Ratio (RCR) Aggregated Score (iCite-RCR-based), described above.

Variables of merit and interest not currently used for ranking:

- OR: median(odds ratio, inverted if <1) supporting gene-trait association.
- N_beta: simple count of beta values with 95% confidence intervals supporting gene-trait association.
- N_snp: SNPs involved with gene-trait association.
- N_study: studies supporting gene-trait association.
- study_N: mean(SAMPLE_SIZE) supporting gene-trait association.
- geneNtrait: total traits associated with the gene.
- traitNgene: total genes associated with the trait.

N_snp, N_study, geneNtrait and traitNgene are counts of the corresponding *unique* entities. From the variables selected via benchmark testing the **meanRankScore** is computed thus:

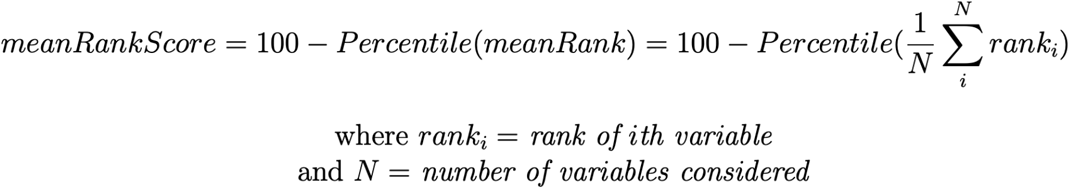

**μ** scores were implemented via the muStat (Wittkowski and Song, 2012) R package. Vectors of ordinal variables represent each case, and non-dominated solutions are cases which are not inferior to any other case at any variable. (For TIGA, cases are genes or traits, corresponding with trait-queries or gene-queries, respectively, and their variables of merit described above.) The set of all non-dominated solutions defines a Pareto-boundary. A **μ** score is defined simply as the number of lower cases minus the number of higher, but the ranking is the useful result. The ranking rule between case k and case k′ may be formalized thus:

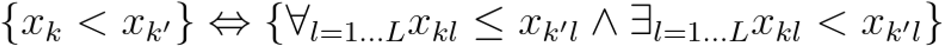

Simply put, case k’ is higher than case k if it is higher in some variable(s) and lower in none.

### Benchmark against gold standard

Lacking a suitable gold standard set of gene–trait associations in general, we instead relied on established gene–disease associations from the Genetics Home Reference, GHR (Fomous *et al*., 2006) and UniProtKB (UniProt Consortium, 2018) databases. This gold stand set was built following a previously described approach (Pletscher-Frankild *et al*., 2015). It consists of 5,366 manually curated associations (positive examples) between 3,495 genes and 709 diseases. All other (2,472,589) possible pairings of these genes and diseases were considered negative examples.

To assess the quality of the TIGA gene–trait associations, we mapped the Ensembl gene IDs to STRING v11 identifiers using the STRING alias file (Szklarczyk *et al*., 2019) and the EFO terms to Disease Ontology identifiers (Schriml *et al*., 2019) based on ontology cross-references and the EMBL-EBI Ontology Xref Service. We then benchmark any individual-variable or multivariate ranking of the associations by constructing the receiver operating characteristic (ROC) curve by counting the agreement with the gold standard.

## Results

### The TIGA web application

TIGA facilitates drug target illumination by currently scoring and ranking associations between protein-coding genes and GWAS traits. While not capturing the entire Catalog, the TIGA app can aggregate and filter GWAS findings for actionable intelligence, e.g., to enrich target prioritization via interactive plots and hitlists (Fig 3), allowing users to identify the strongest associations supported by evidence.

**Fig 3:**
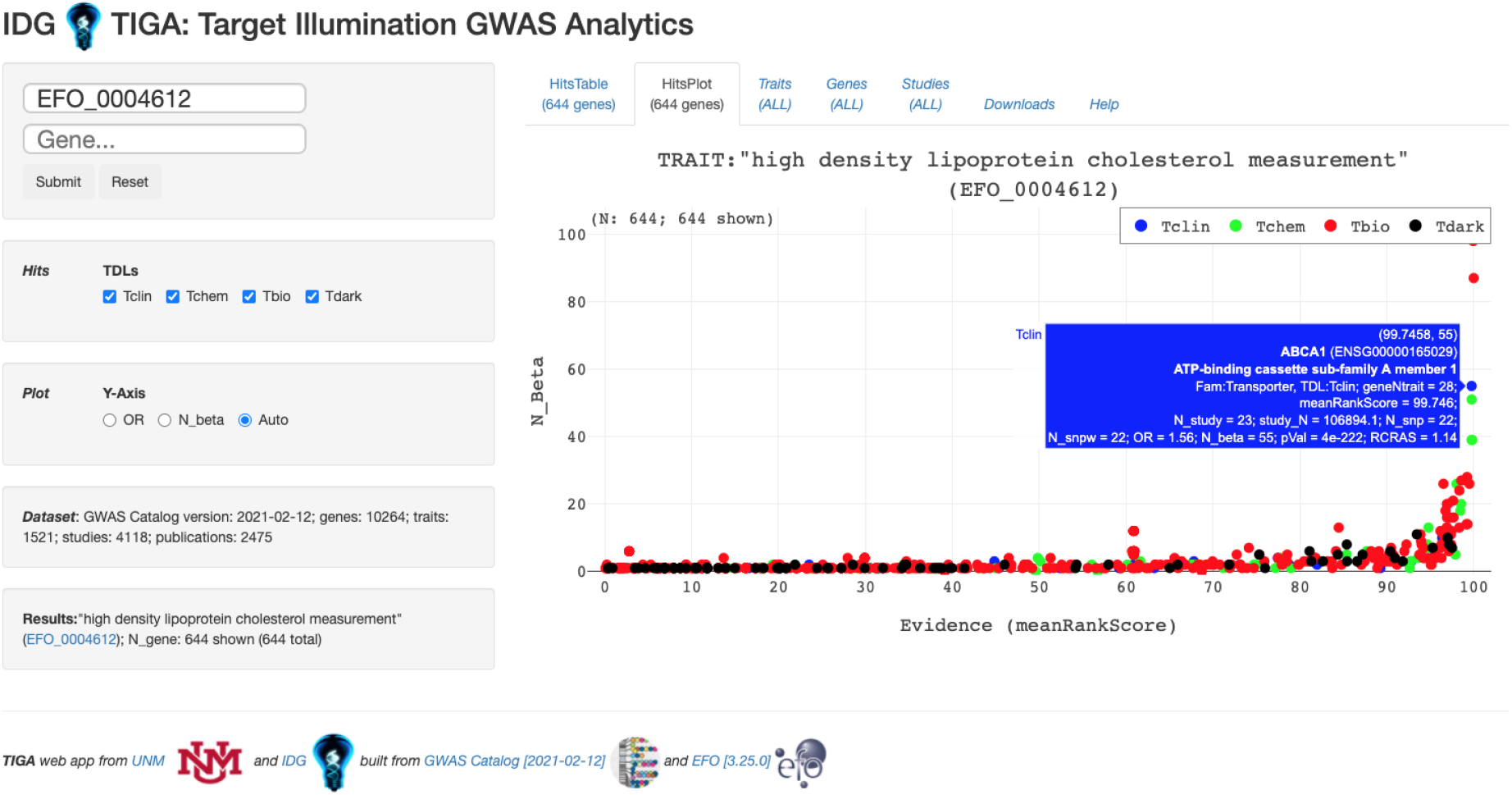
TIGA web application at http://unmtid-shinyapps.net/tiga/, displaying a plot of 644 genes currently associated with trait “high density lipoprotein cholesterol measurement” (EFO_0004612).

Hits are ranked by meanRankScore described in Methods. Scatterplot axes are Effect (OR or N_beta) vs. Evidence as measured by meanRankScore. Plot markers may be sized by pValue or RCRAS. This app accepts “trait” and “gene” query parameters via URL, e.g. ?trait=EFO_0004541, ?gene=ENSG00000075073, ?trait=EFO_0004541&gene=ENSG00000075073. Gene markers are colored by Target Development Level (TDL)(Oprea *et al*., 2018). TDL is a knowledge-based classification that bins human proteins into four categories: **Tclin**, mechanism-of-action designated targets via which approved drugs act (Santos *et al*., 2017; Ursu *et al*., 2019; Avram *et al*., 2020); **Tchem** are proteins known to bind small molecules with high potency; **Tbio** includes proteins that have Gene Ontology (Ashburner *et al*., 2000) “leaf” (lowest level) experimental terms; or meet two of these conditions: A fractional publication count (Pafilis *et al*., 2013) above 5, three or more Gene “Reference Into Function” annotations (Mitchell *et al*., 2003), or 50 or more commercial antibodies in Antibodypedia (Björling and Uhlén, 2008); **Tdark** are manually curated UniProtKB proteins that fail to place in any of the previous categories.

### Benchmark against gold-standard disease–gene associations

To benchmark the quality of the GWAS associations in TIGA, we focused on the 383 EFO terms that could be mapped to diseases and their 20,458 associations with genes. We evaluated the performance of each variable of merit individually against the manually curated gold standard gene–disease associations. The resulting ROC curves showed that the best performing variables are RCRAS, N_study, pVal_mLog, N_snpw, and N_snp, which have areas under the curve (AUC) higher than 0.6 (Fig. 4A and Fig. S.1). The three variables RCRAS, pVal_mLog, and N_snpw are furthermore complementary, having a maximal pairwise Spearman correlation of 0.325, whereas N_study and N_snp are strongly correlated with the better performing RCRAS and N_snpw, respectively. We thus used these three variables as the basis for calculating two multivariate rankings, namely **μ** score and meanRankScore. We benchmarked both rankings the same way as the individual variables and found that **μ** score performs marginally better than meanRankScore score based on their AUC values (Fig. 4B). However, as the meanRankScore outperforms the **μ** score in the area of interest [0.0, 0.2] and is more than five orders of magnitude faster to calculate, we selected it as the final ranking in TIGA. Corresponding plots for the lower performing variables are provided in the supplementary materials for completeness.

**Fig. 4:**
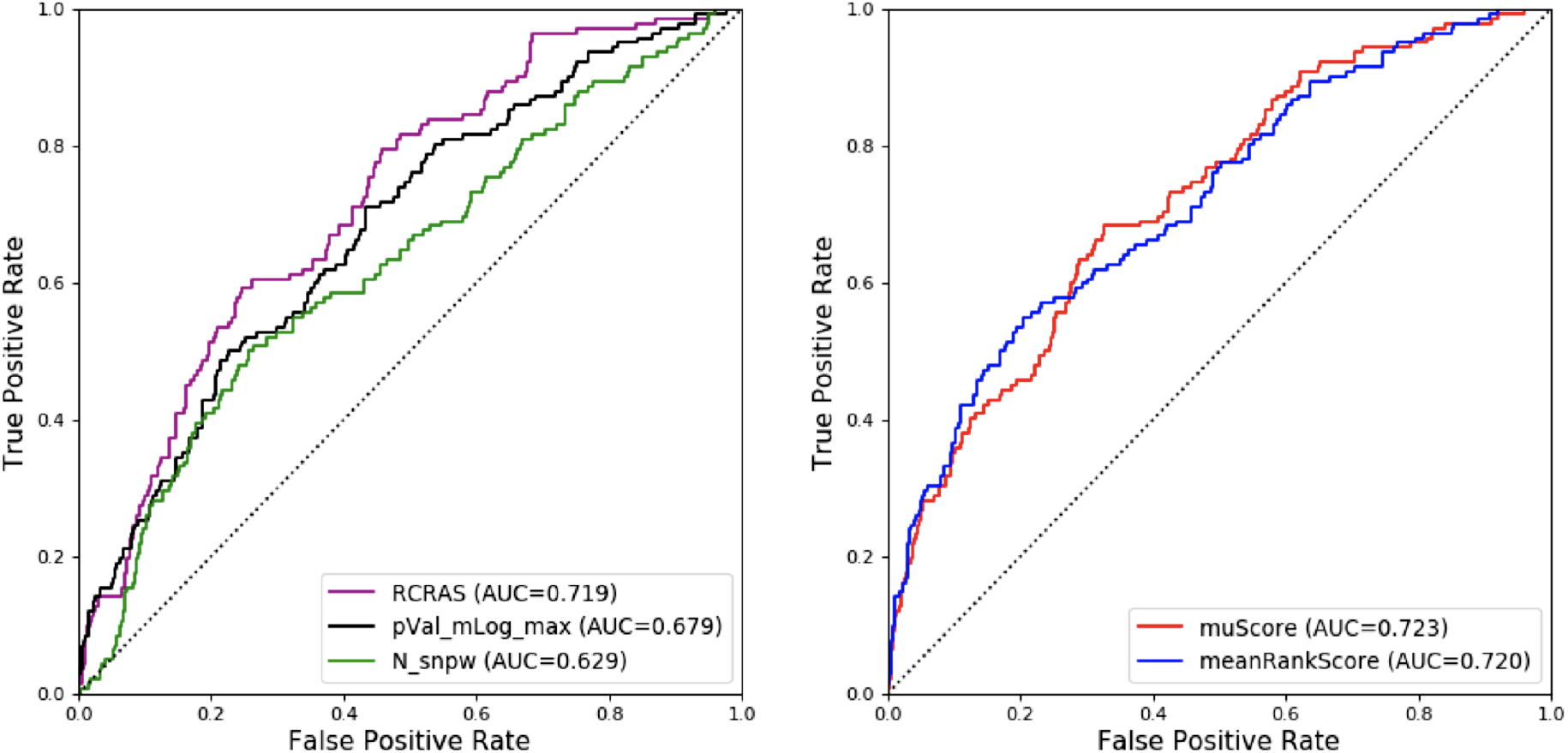
Performance evaluation. The performance of TIGA on the gold standard of gene-disease associations. A) Results for the top-3 individual variables of merit. B) Results for the multivariate ranking by meanRankScore and **μ** score.

### Using TIGA for drug target illumination

The main motivation of developing TIGA is to capture GWAS data when illuminating drug targets. Table 1 shows how many targets from each protein family and IDG Target Development Level (TDL) are covered with associated traits in TIGA, with families as defined by Drug Target Ontology(Lin *et al*., 2017) (DTO) Level 2. IDG TDL is a knowledge based classification: **Tclin** = high-confidence drug targets; **Tchem** = small-molecule modulator exists; **Tbio** = biological function elucidated; **Tdark** = minimal knowledge (Oprea *et al*., 2018). Coverage for the understudied 2,469 **Tdark** proteins is of particular interest. However, the data for other TDLs can also provide unique and complementary evidence, especially in case of **Tbio** proteins that are biologically characterized but have not before been clinically validated.

**Table 1.**
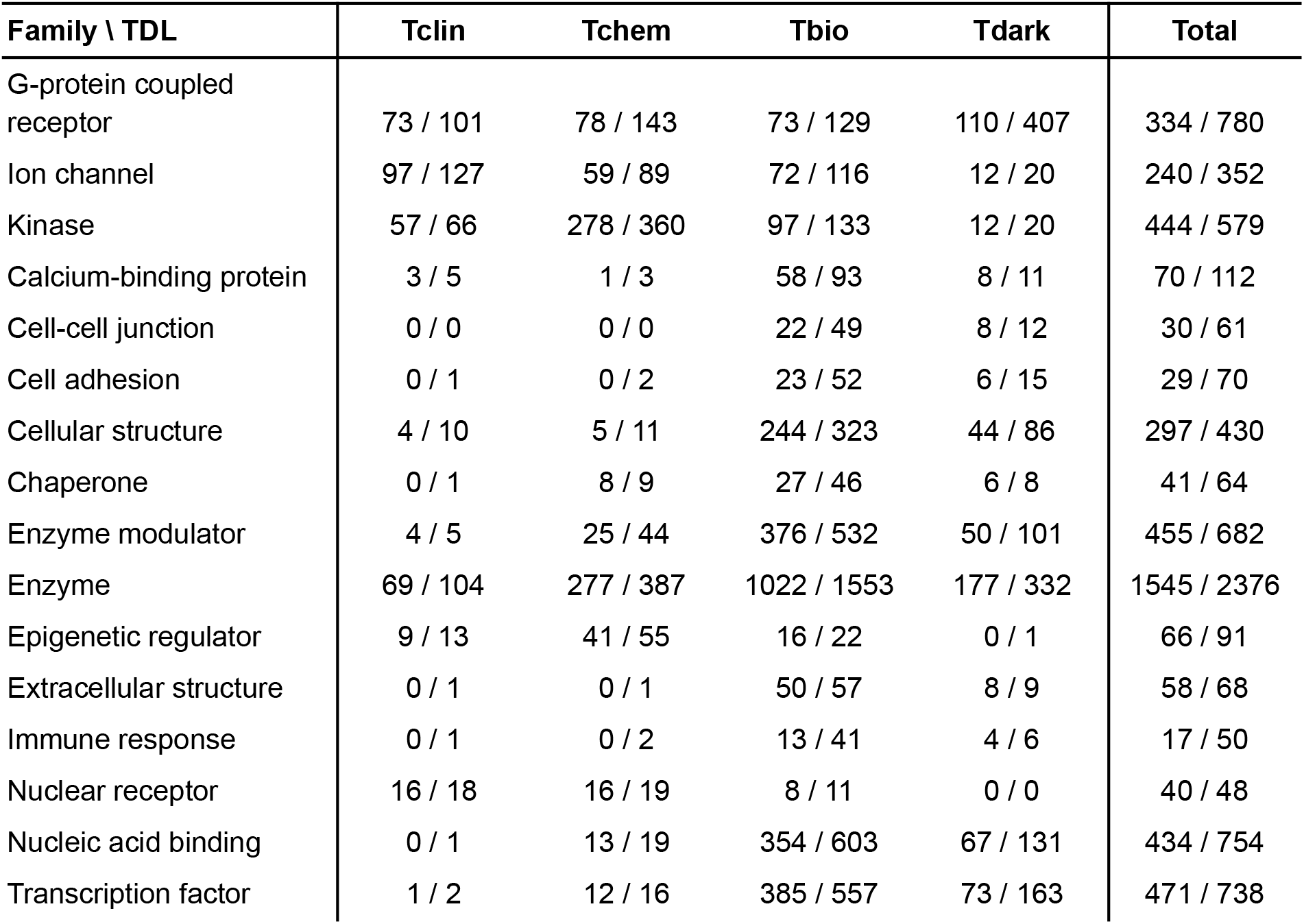
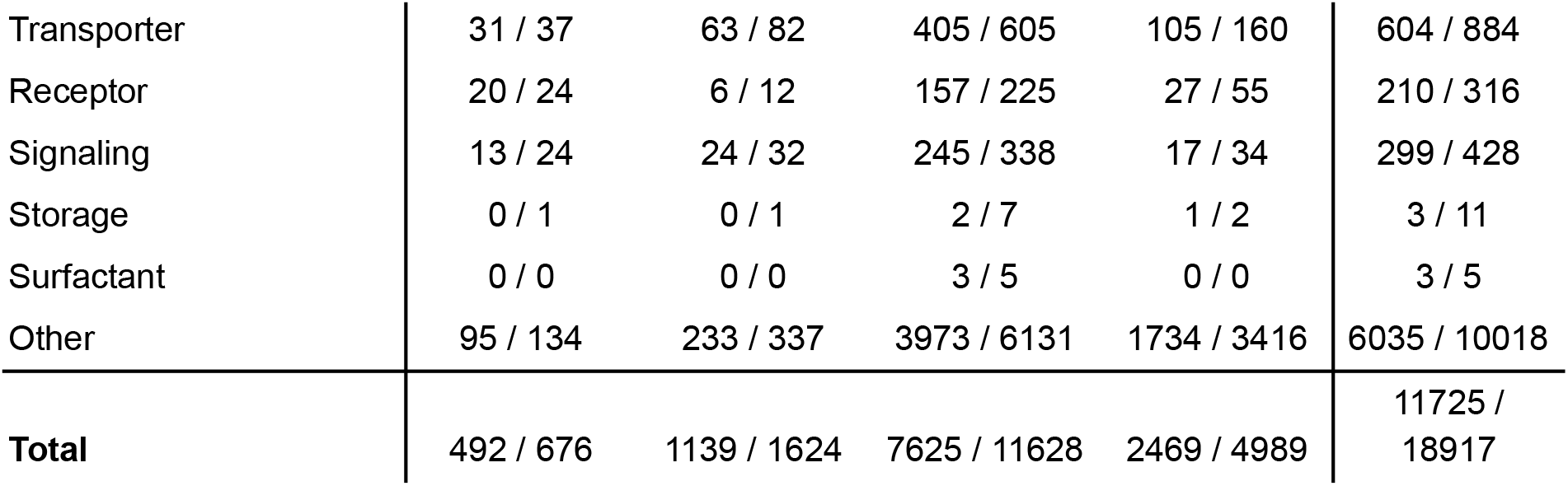
TIGA mapped target (protein) counts by IDG Target Development Level (TDL) and Drug Target Ontology (DTO) level 2 gene family.

Figures 3 and 5 illustrate a typical use case, the plot and gene list for trait “high density lipoprotein cholesterol measurement”, which monitors blood levels of high density lipoprotein cholesterol as a risk factor for heart disease. Figure 6 shows the provenance for one of the associated genes, GIMAP6 “GTPase IMAP family member 6” with the scores and studies for this gene-trait association, including links to the Catalog and PubMed. GIMAP6 is an understudied (**Tbio**) member of the GTPases of immunity-associated protein family (GIMAP). Although literature-based evidence of its potential role in cholesterol homeostasis is scarce (Hoffmann *et al*., 2018; Richardson *et al*., 2020), this finding is substantiated by significantly increased circulating HDL cholesterol levels in GIMAP6-knock-out female mice [https://bit.ly/3uznvCU], suggesting that loss of GIMAP6 function may be associated with hypercholesterolemia-associated disorders.

**Fig. 5:**
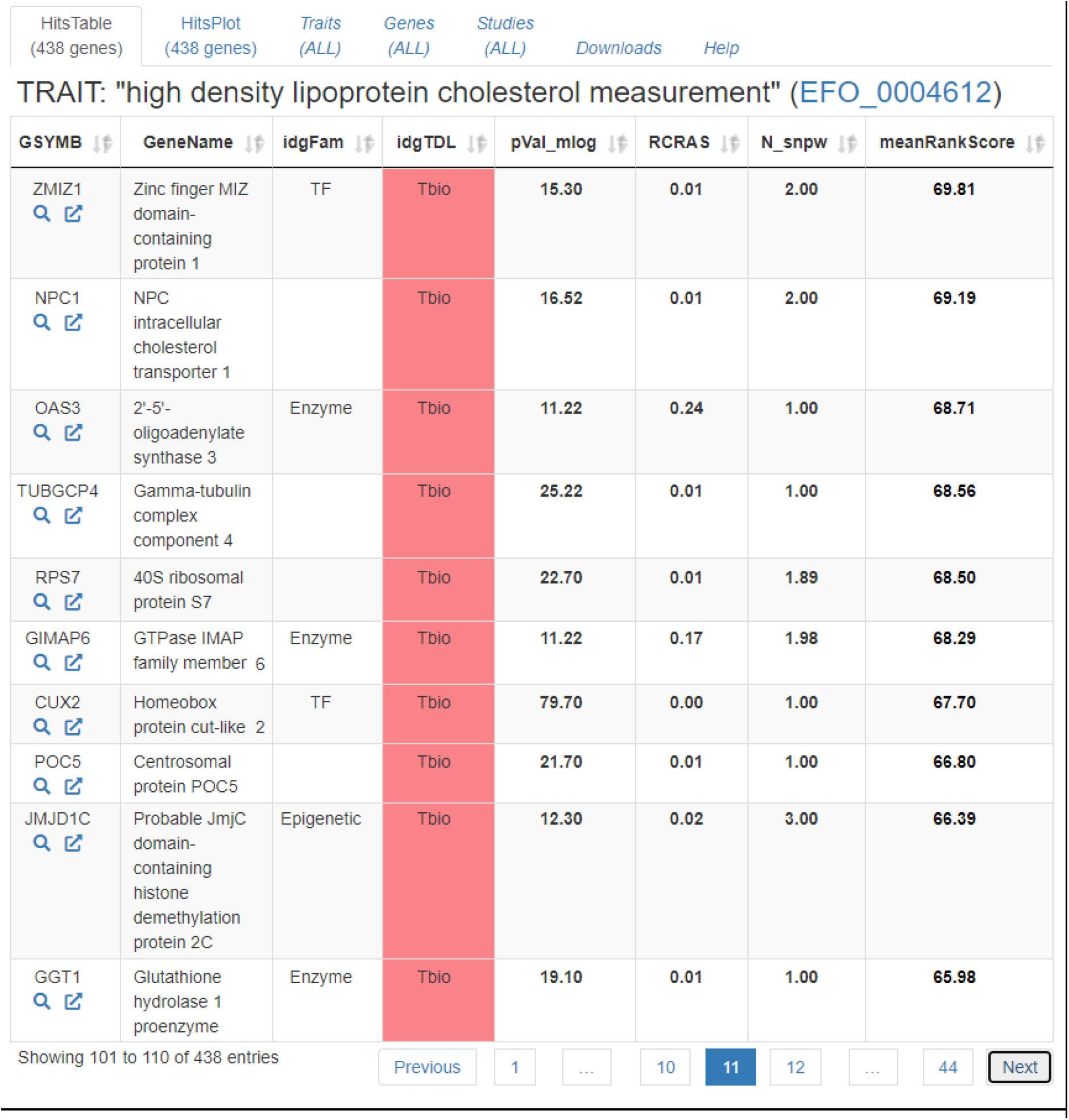
Examples of understudied genes for trait “high density lipoprotein cholesterol measurement” in TIGA.

**Fig. 6:**
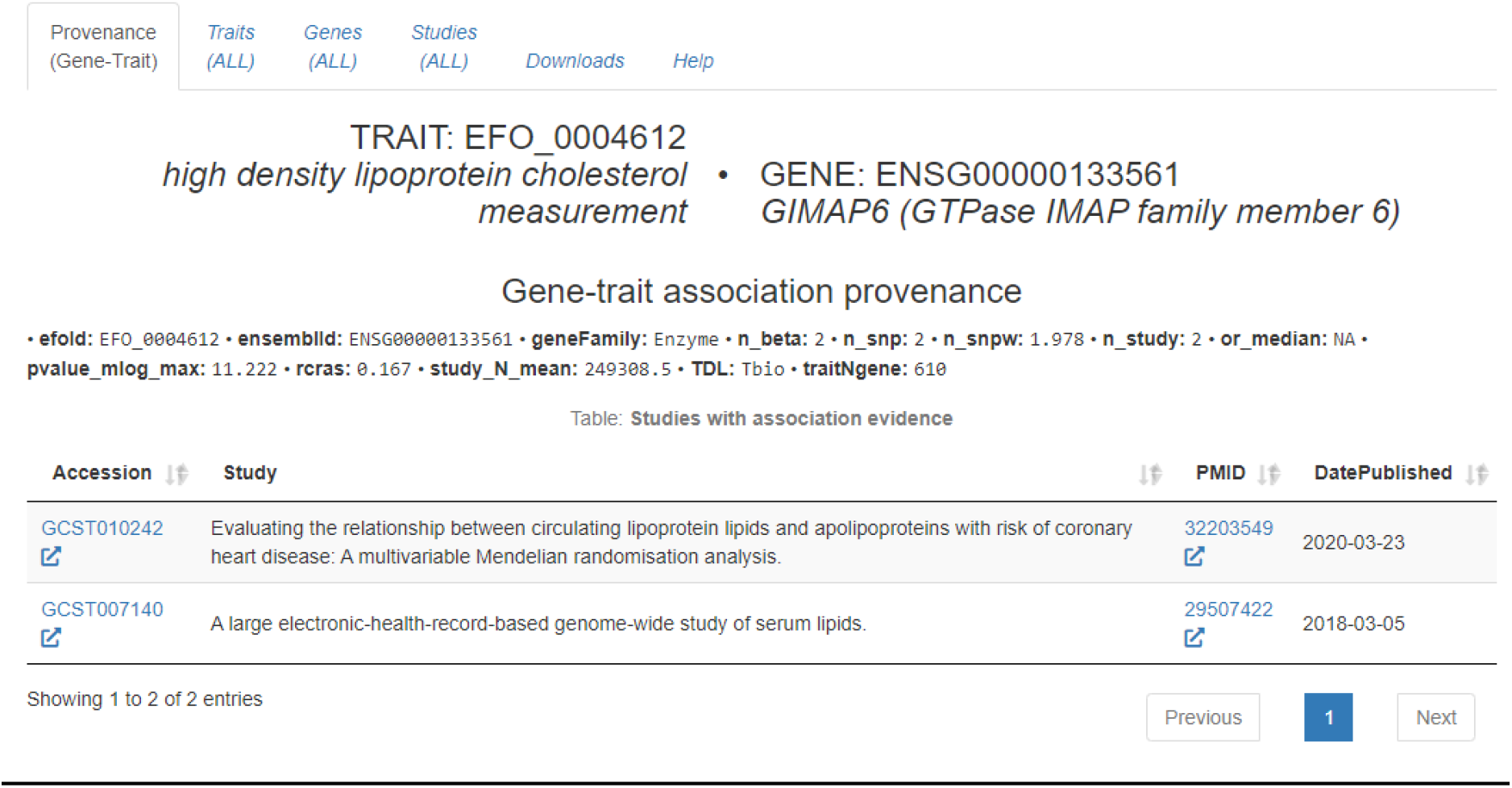
Provenance for association between gene GIMAP6 “GTPase IMAP family member 6” and trait “high density lipoprotein cholesterol measurement”.

Figures 7-8 illustrate another target illumination example, for trait “HbA1c measurement” (glycated hemoglobin, signifying prolonged hyperglycemia), highly relevant to the management of type 2 diabetes mellitus(Rahbar *et al*., 1969; Saudek and Brick, 2009). Figure 8 shows the provenance for one of the associated genes, SLC25A44 “Solute carrier family 25 member 44” with the scores and studies for this gene-trait association, including links to the Catalog and PubMed. SLC25A44 is an understudied (**Tdark**) branched-chain amino acid (BCAA) transporter that acts as metabolic filter in brown adipose tissue, contributing to metabolic health (Yoneshiro *et al*., 2019). SLC25A44 may be involved in subcutaneous white adipose BCAA uptake and catabolism(Lee *et al*., 2021).

**Fig 7:**
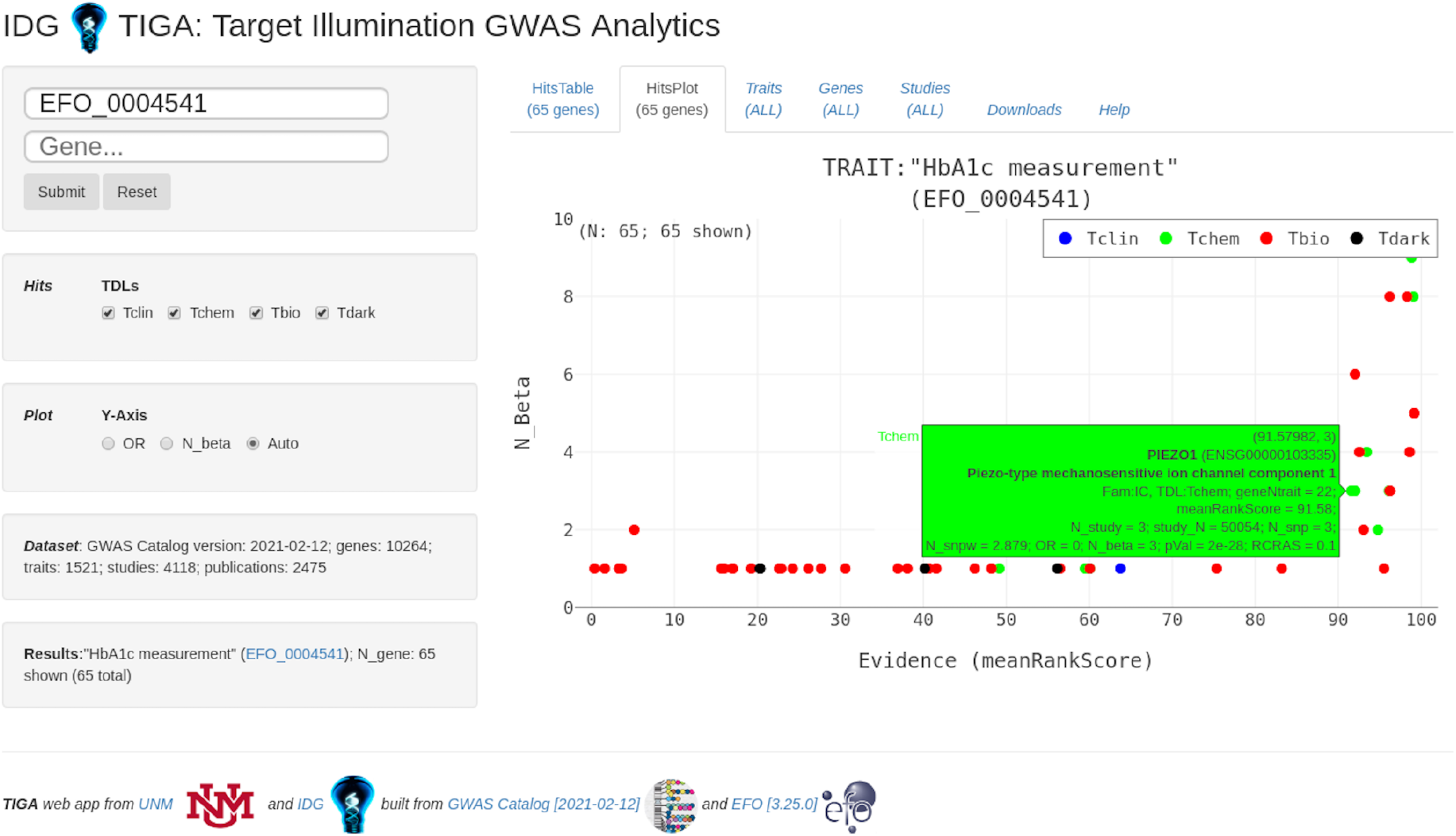
TIGA web application at http://unmtid-shinyapps.net/tiga/, displaying a plot of genes associated with trait “HbA1c measurement” (EFO_0004541).

**Fig. 8:**
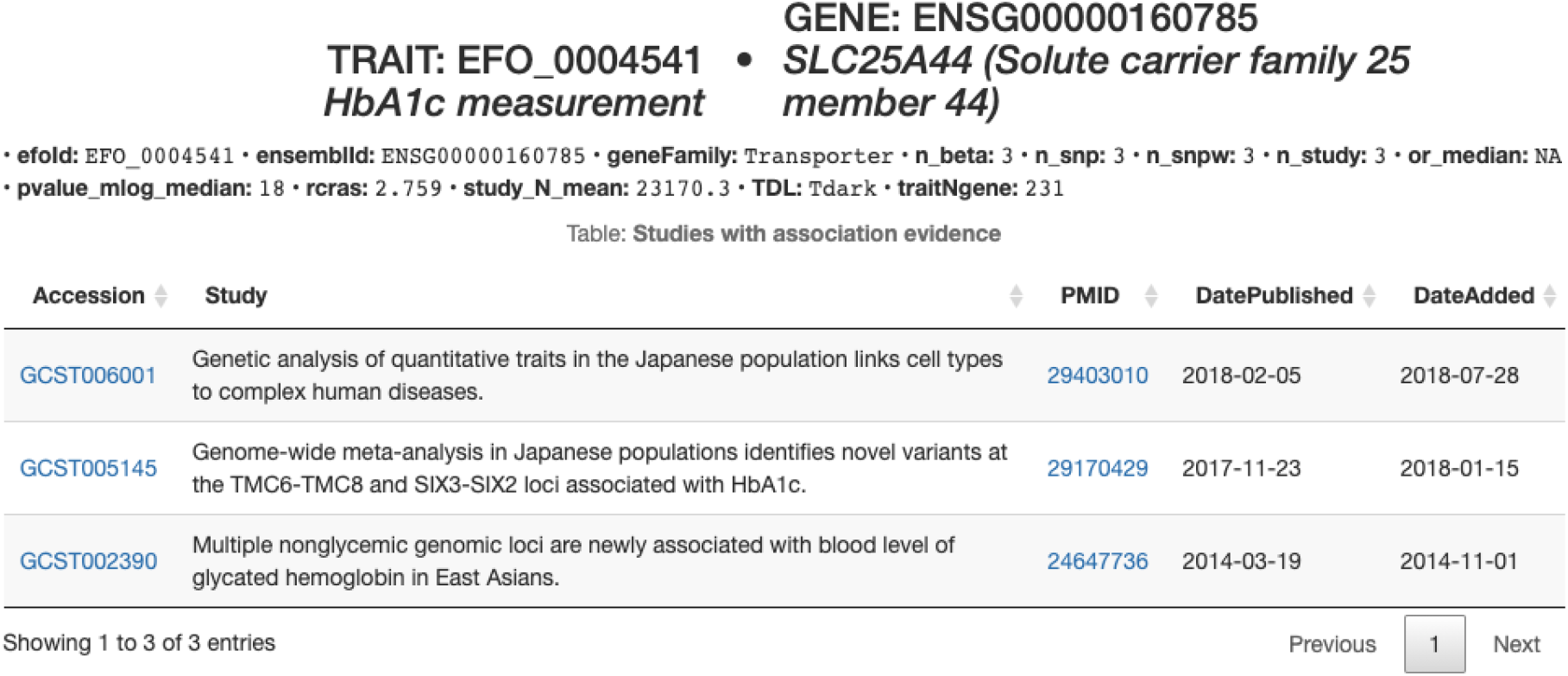
Provenance for association between gene SLC25A44 “Solute carrier family 25 member 44” and trait “HbA1c measurement”.

## Discussion

### Target illumination

The explicit goal of the NIH Illuminating the Druggable Genome (IDG) program (Oprea *et al*., 2018) is to “map the knowledge gaps around proteins encoded by the human genome.” TIGA is fully aligned with this goal, as it evaluates the GWAS evidence for disease (trait) – gene associations. TIGA generates GWAS-centric trait–gene association dataset using an automated, sustainable workflow amenable for integration into the Pharos portal (Nguyen *et al*., 2017; Sheils *et al*., 2021). The Open Targets platform (Ochoa *et al*., 2020; Ghoussaini *et al*., 2020) uses Catalog data and other sources, assisted by supervised machine learning, to identify probable causal genes, and validate therapeutic targets by aggregating and scoring disease–gene associations for “practicing biological scientists in the pharmaceutical industry and in academia.” Open Targets associations are enhanced, but also limited, by the training data and knowledge sources reflecting current understandings of genetics. In contrast, TIGA provides aggregated evidence solely from the Catalog, reflecting experimental results with minimal or no bias, thus more interpretable in terms of provenance and methodology, and more suitable for some downstream consumers and use cases.

### From information to useful knowledge

In data-intensive fields such as genomics, specialized tools facilitate knowledge discovery, yet interpretation and integration can be problematic for non-specialists. Accordingly, this unmet need for integration and interpretation requires certain layers of abstraction and aggregation, which depend on specific use cases and objectives. Our target audience is drug discovery scientists for whom the aggregated findings of GWAS, appropriately interpreted, can provide additional value as they seek to prioritize targets. This clear purpose serves to focus and simplify all aspects of its design. Our approach for evidence aggregation is simple, easily comprehensible, and based on what may be regarded as axiomatic in science and rational inductive learning: First and foremost, evidence is measured by counting independent confirmatory results.

Interpretability concerns exist throughout science, but GWAS is understood to present particular challenges (Lambert and Black, 2012; Visscher *et al*., 2017; Marigorta *et al*., 2018; Gallagher and Chen-Plotkin, 2018). The main premise of GWAS is that genotype-phenotype correlations reveal underlying molecular mechanisms. While correlation does not imply causation, it contributes to plausibility of causation. Genomic dataset size adds difficulty.

The standard GWAS p-value significance threshold is 5e-8, based on overall p-value 0.05 and Bonferroni multiple testing adjustment for 1-10 million tests/SNPs (Marigorta *et al*., 2018). The statistical interpretation is that the family-wise error rate (FWER), or overall probability of a type-1 error, is 5%, but associations to mapped genes require additional interpretation. Motivated by, and despite these difficulties, it is our belief that GWAS data can be rationally interpreted and used by non-specialists, if suitably aggregated. Accordingly, TIGA is a rational way to suggest and rank research hypotheses, with the caveat that the identified signals may be accompanied by experimental noise and systematic uncertainty.

### Designing for downstream integration

Biomedical knowledge discovery depends on integration of sources and data types which are heterogeneous in the extreme, reflecting the underlying complexity of biomedical science. These challenges are increasingly understood and addressed by improving data science methodology. However, provenance, interpretability and confidence aspects are underappreciated and rarely discussed. As in all signal propagation, errors and uncertainty accrue and confidence decays. Here, we proposed the use of simple, transparent, and comprehensible metrics to assess the relative confidence of disease-gene associations, via the unbiased meanRank scores. Figure 9, summarizing TIGA sources and interfaces, illustrates its well-defined role. Continuous confidence scores support algorithmic weighting and filtering. Standard identifiers and semantics support rigorous integration. Limiting provenance to the Catalog and its linked publications, semantic interpretability is enhanced.

**Fig 9:**
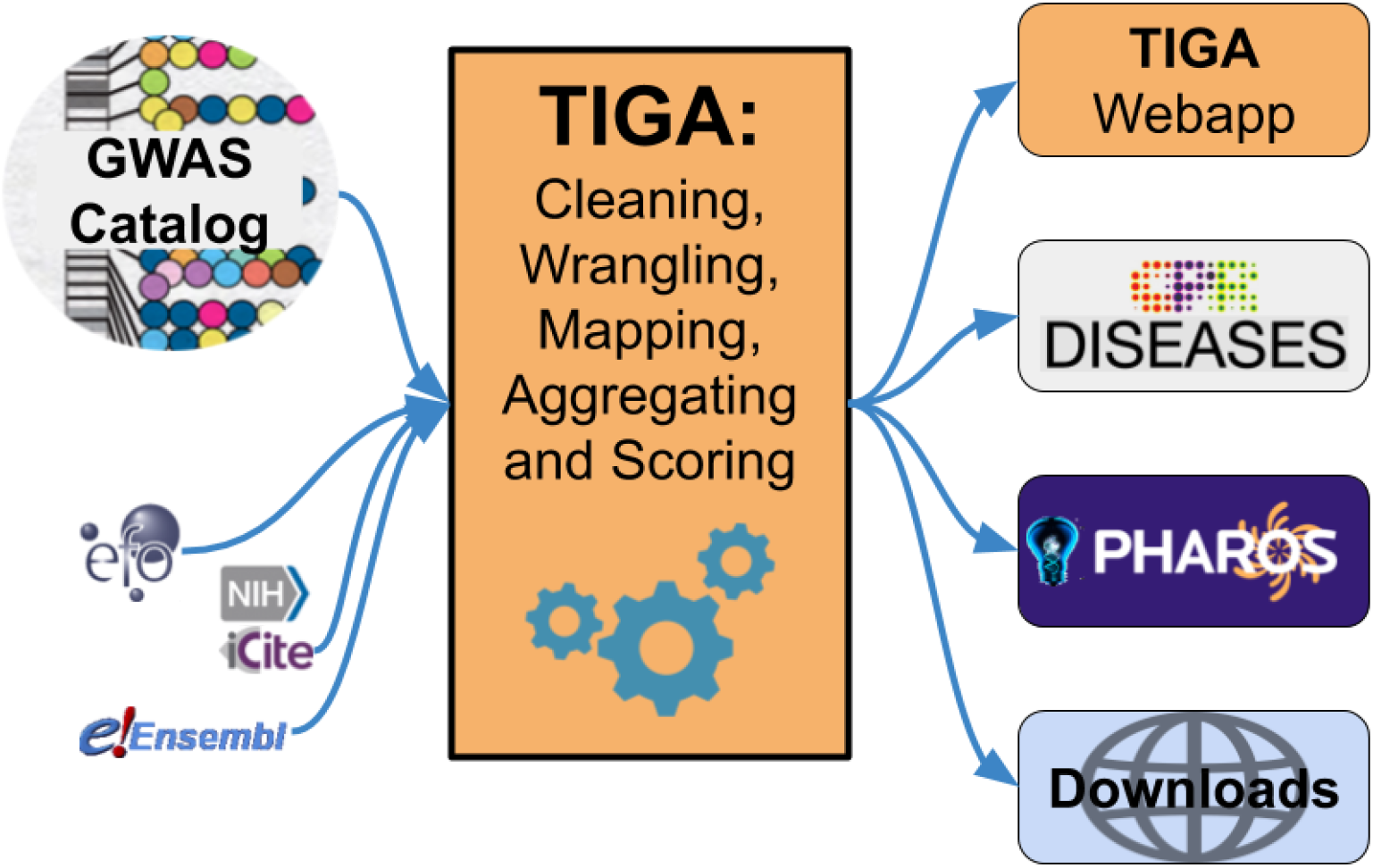
TIGA data sources and interfaces. TIGA integrates GWAS data from the Catalog and several other sources to rank gene-disease associations. These associations can be accessed through the TIGA webapp and are integrated into the DISEASES (Pletscher-Frankild *et al*., 2015) and Pharos platforms. Bulk download is also available.

## Conclusions

We agree with Visscher et al. that: “the paradigm of ‘one gene, one function, one trait’ is the wrong way to view genetic variation”(Visscher *et al*., 2017). Yet given the intrinsic complexity of biomedical science, progress often requires simplifying assumptions. Findings must be interpreted in context for an audience and application. Mindful of these concerns and limitations, TIGA provides a directly interpretable window into GWAS data, specifically for drug target hypothesis generation and elucidation. As interest in “interpretable machine learning” and “explainable artificial intelligence” (Gilpin *et al*., 2018) grows, TIGA summarizes gene-trait associations derived solely and transparently from GWAS summary- and meta-data, with rational and intuitive evidence metrics and a robust, open-source pipeline designed for continual updates and improvements. Whether in stand-alone mode, or by integration with other interfaces, TIGA aims to contribute to drug target identification and prioritization.

## Acknowledgements

This work was supported by US National Institutes of Health grant U24 224370 for “Illuminating the Druggable Genome Knowledge Management Center” (IDG KMC), and by the Novo Nordisk Foundation (grant number NNF14CC0001).

## Abbreviations and Definitions

Common terms used in GWAS and related fields can vary in their definitions and connotations depending on context. Therefore for clarity and rigor the following definitions are provided, which we consider consistent with best practices in the GWAS and drug discovery communities.

|  |  |
| --- | --- |
| **genotype** | An organism has one genotype, comprised of a germ line genome and multiple somatic genomes. Statistical models may assume a population distribution hence a population genotype. |
| **phenotype** | An organism has one phenotype, comprised of (potentially) all non-genomic observable characteristics, a.k.a. phenotypic traits. |
| **gene** | Genomic unit responsible for an expression product. Protein coding genes are a subset of this definition. |
| **trait** | Single non-genomic, observable characteristic. |
| **drug target** | Biomolecular entity involved in the mechanism of action of a drug. The IDG project is protein-centric; hence drug targets are all proteins. |

## Supplementary material

### Benchmark using additional variables against gold-standard disease–gene associations

**Supplementary Fig 1:**
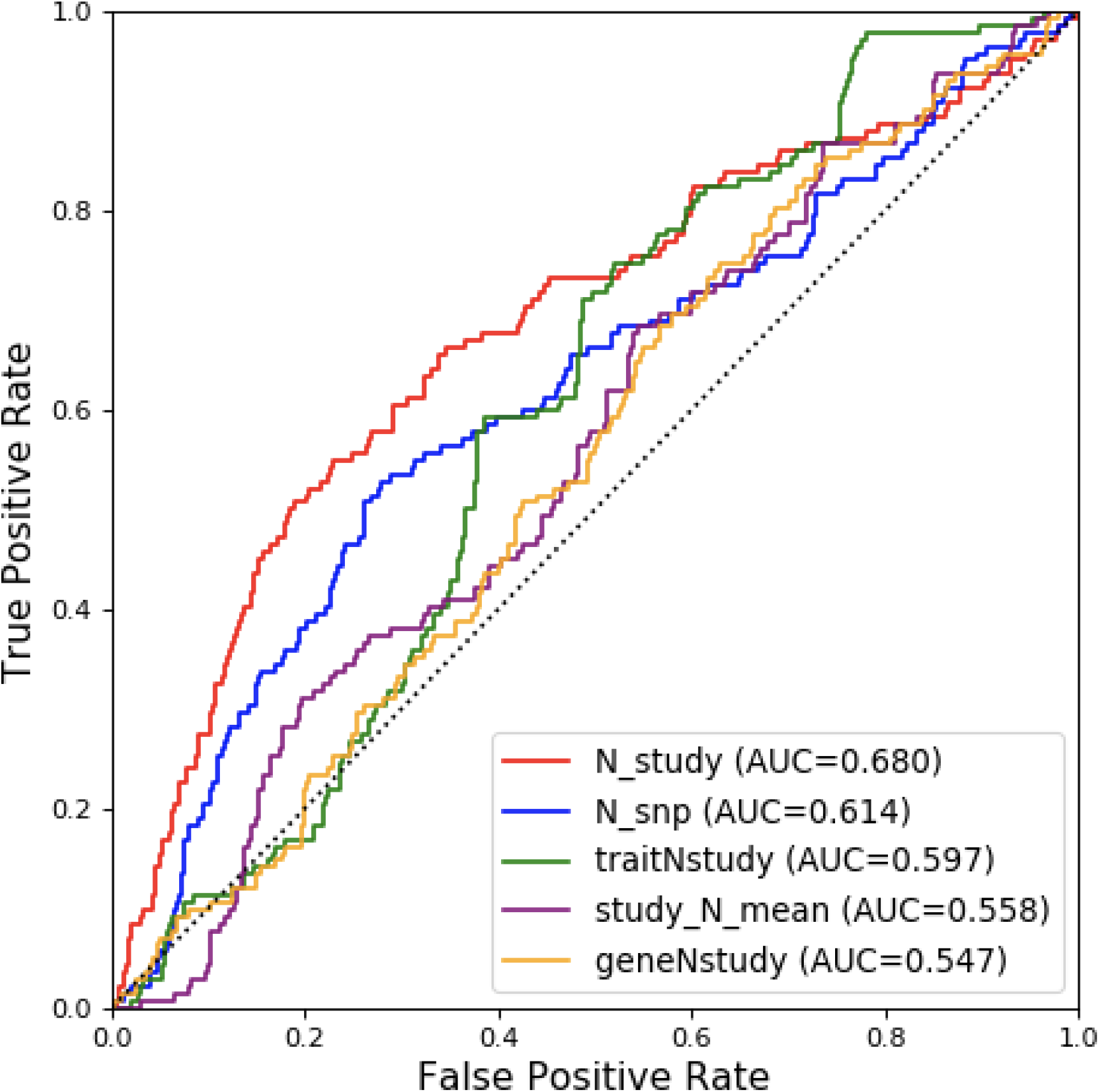
The performance of TIGA variables on the gold standard of gene-disease associations. Results for additional individual variables of merit ***not*** in the top 3 selected for TIGA scoring and included in Fig 4.

**Supplementary Fig 2:**
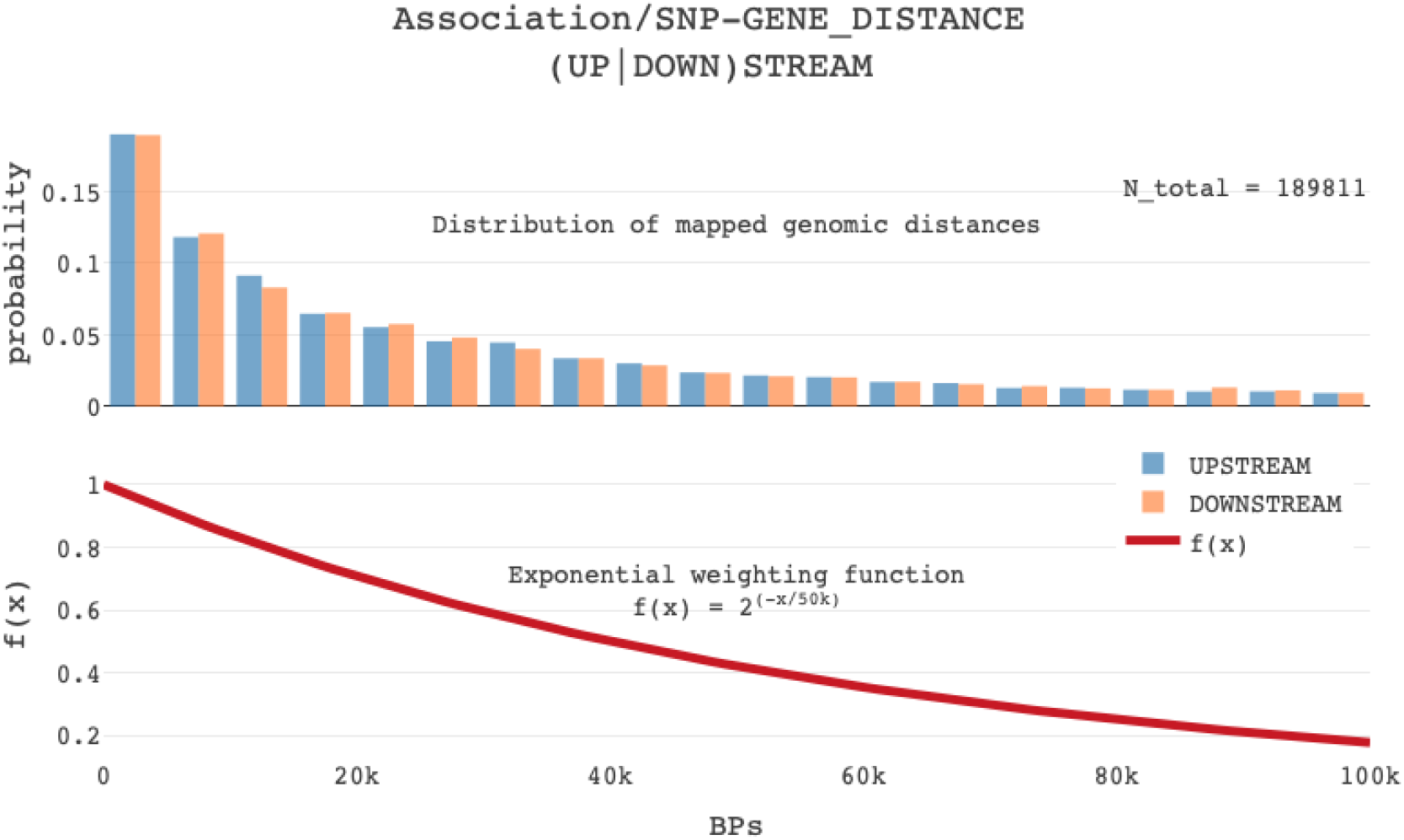
SNP-gene distances for (up|down)stream genes and TIGA weighting function

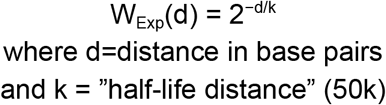

## Footnotes

1 GWAS Catalog FAQ, https://www.ebi.ac.uk/gwas/docs/faq, accessed 24 Sept 2019.

